# Negative Life Events and Epigenetic Ageing: a Study in the Netherlands Twin Register

**DOI:** 10.1101/2024.02.20.581138

**Authors:** B.M.A. Gonggrijp, S.G.A. van de Weijer, C.C.J.H. Bijleveld, D.I. Boomsma, J. van Dongen

## Abstract

We aimed to understand the long-term impact of negative life events (NLE) on epigenetic aging in 1,808 adults from the Netherlands Twin Register, analyzing five epigenetic biomarkers (Hannum, Horvath, PhenoAge, GrimAge, DunedinPACE) and a series of NLE, including victimization and economic hardship. In population-level analyses, associations between a higher number of NLE (particularly financial adversities, sexual crimes, and job loss) were seen for GrimAge and DunedinPACE biomarkers. The association between the number of NLE and financial problems and epigenetic age acceleration measured by the GrimAge biomarker persisted after adjusting for BMI, smoking, and white blood cell counts. In monozygotic twin pairs discordant for NLE (274 pairs) the associations were diminished, indicating that the population associations may be confounded by shared familial (genetic and environmental) factors. These findings underscore the intricate link between environmental stressors and biological aging, stressing the need for comprehensive studies considering both genetic and environmental influences.

---

Negative life events influence numerous aspects of lives of individuals who experience these events. Extensive prior research has consistently linked the experience of negative life events to negative life outcomes across domains such as relationships, employment, and elevated risks of health problems, including mental health problems, poor cognitive functioning, cardiovascular disease, gastrointestinal complications, musculoskeletal problems, reproductive challenges, compromised immune function, and disruptions in the endocrine system (e.g., Dag et al., 2020; Monaghan & Haussmann, 2015; Patton et al., 2003; Young & Schieman, 2012). Previous research also revealed a connection between negative life events and premature mortality. For example, Pridemore and Berg (2017) found that men who had recently experienced violence have 2.6 times higher odds of facing premature death in comparison to those who had not experienced this. Research into adverse childhood experiences has revealed profound and enduring effects on lifespan: a study by Brown et al. (2009) found that individuals who have endured six or more of such experiences typically have a life expectancy nearly 20 years shorter than those without such adversities.

Given the substantial evidence linking negative life events to a range of poor life outcomes and health, we ask how we may understand the underlying biological mechanisms that mediate these effects. While multiple factors contribute to the complex interplay between exposure to negative life events and health outcomes, one area gathering significant attention is the role of epigenetics. Epigenetic changes to the DNA code affect gene expression, without changing the code itself. In large-scale epidemiological studies, epigenetic changes can be measured by differential DNA methylation. DNA methylation is a process where a methyl group is added to DNA, altering gene activity without changing the DNA sequence itself, and influencing the way genes are expressed. Epigenetic markers can be assessed on arrays that assess 400.000 to ∼900.000 markers. Based on such DNA methylation data, multiple markers of biological aging have been proposed. These biological markers are developed through algorithms that estimate an individual’s ‘DNA methylation age’ based on the methylation levels at specific points in the DNA, known as CpG sites (Raffington & Belsky, 2022). The deviation of DNA methylation age from chronological age, referred to as “epigenetic age acceleration”, has been linked to increased mortality risk (Marioni et al., 2015a) and a range of adverse health outcomes, such as cognitive impairment and poor physical and cognitive fitness (Levine, Bennet & Horvath, 2015; Marioni et al., 2015b). The first generation of these epigenetic age acceleration measures included the Hannum and Horvath ‘clocks’ and were designed to compare the biological age of older individuals with that of younger ones. These early epigenetic biomarkers utilized machine learning to estimate a person’s chronological age based on DNA samples collected at a specific ages. While these first-generation biomarkers showed moderate effectiveness in predicting mortality, they predicted less well when forecasting other age-related conditions, including diseases, disability, and declines in physiological and functional capacities (Horvath, 2013; Hannum et al., 2013; Bell et al., 2019). The development of next-generation epigenetic markers like PhenoAge and GrimAge is a two-step process aimed at predicting an individual’s lifespan using survival analysis. Initially, PhenoAge is created by correlating physiological markers with remaining lifespan, generating a score based on these markers. The process then extends to incorporate DNA methylation data, refining the PhenoAge score to a DNA methylation-based version. GrimAge, while also employing machine learning, further includes factors like age, sex, and smoking history, offering a more comprehensive assessment of physiological aging. Both biomarkers have proven to be more accurate in predicting morbidity and mortality compared to the first-generation biomarkers (Levine et al., 2018). Recently, a pace of aging measure has emerged, the DunedinPACE. DunedinPACE evaluates the rate of aging by analyzing changes over time in multiple physiological systems. This algorithm was trained to integrate information from a range of biomarkers across different body systems, like cardiovascular, metabolic, renal, immune, dental, and pulmonary functions. By assessing these changes over time, DunedinPACE can estimate the rate at which these systems are aging in each individual. Thus, while the Hannum, Horvath, PhenoAge, and GrimAge are interpreted as biological age in years, DunedinPACE values represent rates of aging (Belsky et al., 2022; Raffington & Belsky, 2022). Table 1 provides an overview of the epigenetic biomarkers we will consider, with the criterion used, information about the discovery sample, and interpretation of the measure’s values.

**Table 1.** Description of three generations of biological aging measurements.

| Measure | Training Tissue | Training Sample | Age | Predicted trait |
| --- | --- | --- | --- | --- |
| **<u>First Generation: Chronological age predictors</u>** |  |  |  |  |
| Horvath | 51 healthy tissues and cell types | 7844 Adults across 82 different datasets across entire lifespan | 0-100 | Chronological age predicted by DNA methylation |
| Hannum | Whole blood | 482 adult volunteers at UC San Diego, University of Southern California, and West China Hospital. | 19-101 |  |
| **<u>Second Generation: Mortality risk predictors</u>** |  |  |  |  |
| PhenoAge | Whole blood | 9926 adults from the InCHIANTI Study | 21-100 | Blood-chemistry PhenoAge |
| GrimAge | Whole blood | 6935 Adults from the Framingham Heart Study Offspring Cohort. | 53-73 | Mortality risk |
| **<u>Third generation: Pace of Aging</u>** |  |  |  |  |
| DunedinPACE | Whole Blood | 1037 participants from the Dunedin Study 1972 - 1973 birth cohort. | Measured at 26, 32, 38 and 45 years | Physiological decline experienced per 1 year of calendar time. |

A handful of studies examined the relationship between exposure to adverse childhood life events and biological aging. For instance, Sumner et al. (2023) collected saliva samples for DNA isolation in 161 participants aged 8 to 16 years and discovered that experiencing everyday stressful life events during adolescence was associated with accelerated epigenetic aging as indexed by the Horvath measure. Joshi, Lin, and Raina (2023) conducted a study involving 1445 individuals aged 45 to 85 from whom blood samples were collected. They explored second-generation measures and found that childhood exposure to parental separation or divorce and emotional abuse was linked to higher GrimAge acceleration but not to PhenoAge acceleration in later life. Raffington et al. (2021) examined 600 children and adolescents aged 8 to 18 in the Texas Twin Project, measuring saliva DNA methylation and socioeconomic circumstances. Children from more disadvantaged families and neighborhoods exhibited a faster pace of aging as measured by DunedinPACE. Kim et al. (2023) examined 895 adults at age 15 and later followed up with 867 of these individuals at age 20, conducting DNA methylation profiling in blood samples. Their research revealed that individuals who had experienced a higher number of adverse childhood experiences displayed accelerated biological aging for the PhenoAge, GrimAge, and DunedinPACE. This effect persisted into midlife and earlier adulthood, even after accounting for demographic, behavioral, and socioeconomic variables. Research on experiencing negative life events during adulthood has focused on Post-Traumatic Stress Disorder (PTSD), with conflicting findings. For instance, Wolf et al. (2019) reported that PTSD was linked to faster aging based on the Horvath but not the Hannum measure, while Oblak et al. (2021) found an age acceleration for the Hannum, but deceleration for the Horvath measure. Wang et al. (2022) conducted a cross-sectional co-twin control study design with DZ and MZ twins discordant for current PTSD to control for shared genetic and other familial factors. They found that twins with PTSD exhibited significantly advanced DNA methylation age acceleration compared to their twin brothers without PTSD across several biomarkers — Horvath, Hannum, and PhenoAge — but not GrimAge. These findings indicated that, across most measures of DNA methylation age acceleration, twins with current PTSD were ‘epigenetically older’ by an estimated 1.6 to 2.7 years of biological age than their unaffected twin siblings and suggest that the effect is not influenced by genetic or shared environmental confounding. In contrast, Yang et al. (2020) and Katrinli et al. (2020) found an association of PTSD with GrimAge acceleration, underscoring the varying impacts of PTSD on different biological aging markers. Thus, while adverse childhood experiences are linked to acceleration in the Horvath, PhenoAge, GrimAge, and DunedinPACE measures, findings regarding traumatic stress disorder during adulthood and accelerated aging are less consistent.

We aim to contribute to the existing literature by investigating the association between negative life events and blood-cell-derived epigenetic age acceleration in adulthood in a large population-based sample of twin families from the Netherlands Twin Register (N=1808, including 421 monozygotic twin pairs). Most previous research has centered primarily on childhood adversity or traumatic events in the context of PTSD, or focused on a restricted subset of epigenetic biomarkers. Our study enriches the existing literature by examining a broad spectrum of negative life events, ranging from crime victimization and financial troubles to losing a loved one in a population-based cohort of adults, and look at the five most widely researched biomarkers: Hannum, Horvath, PhenoAge, GrimAge, and DunedinPACE. In our study, we initially explored the link between the total number of negative life events and epigenetic age acceleration in all participants. Next, we investigated the association between specific life events and epigenetic age acceleration. Finally, we conducted within-pair analyses among monozygotic twins who were discordant for life events. This part of the study focused on comparing epigenetic age acceleration between twins who experienced a different number of negative life events. This analysis controls for shared environmental and genetic confounding (Gesell, 1942; Gonggrijp et al., 2023; Wang et al., 2022) and aims to elucidate the direct impact of negative life events on epigenetic aging, independent of genetic and shared environmental factors. To the best of our knowledge, this research is the first to implement a discordant twin analysis concerning epigenetic biomarkers and negative life events. By examining the relationship between epigenetic age and negative life events, this research seeks to shed light on the underlying mechanisms linking negative life events to negative health outcomes.

## Methods

### Cohort description

Data were collected from participants from the Netherlands Twin Register (NTR). Respondents fill out surveys on health and lifestyle every two to three years. Full details about data collection have been reported previously (e.g. Ligthart et al. 2019; Distel et al. 2011). DNA was collected from buccal cells and whole blood as part of multiple projects. For the current paper, we analyzed DNA methylation measured in whole blood collected in the NTR-Biobank study, conducted in 2004–2008 and 2010–2011, (van Dongen et al., 2016; Willemsen et al., 2010). Good quality whole blood DNA methylation data were available from twins, parents of twins, siblings of twins, and spouses of twins. Data regarding life events were measured in a survey from 2009. Respondents were only included in the sample if the survey was administered prior to the DNA sample collection. The selection of participants is detailed in Figure 1, resulting in a total sample size of 1,808 individuals with data on life events and epigenetic biomarkers.

**Figure 1.**
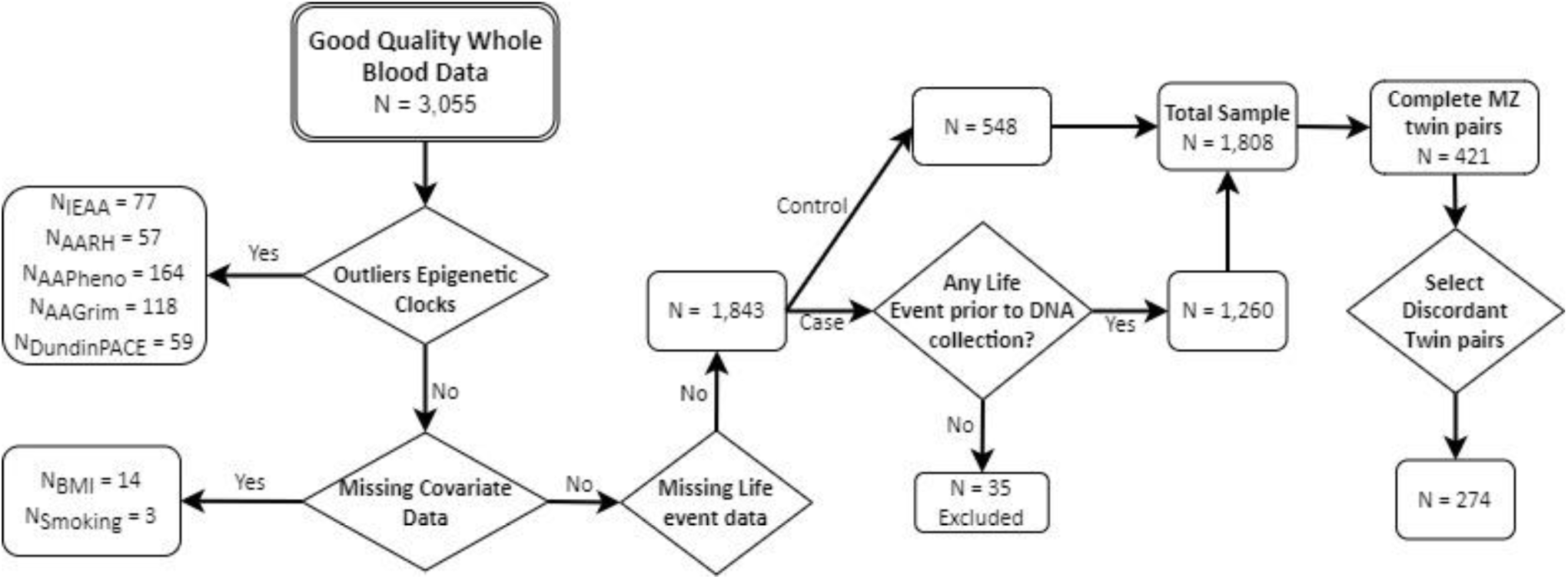
Flowchart of Data Selection. Note: AARH = Hannum Age Acceleration, IEAA = Intrinsic Epigenetic Age Acceleration, AAPheno = PhenoAge Acceleration, AAGrim = GrimAge acceleration, WB = Whole Blood. Control depict participants who did not experience any life events.

### Ethics

Written informed consent was obtained from all participants. The study was approved by the Central Ethics Committee on Research Involving Human Subjects of the VU University Medical Centre, Amsterdam, an Institutional Review Board certified by the U.S. Office of Human Research Protections (IRB number IRB00002991 under Federal-wide Assurance-FWA00017598; IRB/institute codes, NTR 03-180).

### DNA Collection

#### DNA Methylation

DNA methylation was assessed with the Infinium HumanMethylation450 BeadChip Kit (Illumina, San Diego, CA, USA) by the Human Genotyping facility (HugeF) of ErasmusMC, the Netherlands (http://www.glimdna.org/) as part of the Biobank-based Integrative Omics Study (BIOS) consortium (Bonder et al., 2017). DNA methylation measurements have been described previously (Bonder et al., 2017; van Dongen et al., 2016) Genomic DNA (500ng) from whole blood underwent bisulfite treatment with the Zymo EZ DNA Methylation kit (Zymo Research Corp, Irvine, CA, USA), and 4 µl of bisulfite-converted DNA was measured on the Illumina 450k array following the manufacturer’s protocol. A number of sample- and probe-level quality checks and sample identity checks were performed, as described in detail previously (van Dongen et al., 2016). In short, sample-level QC was conducted with the assistance of MethylAid (van Iterson et al., 2014). Probes were set to missing in a sample if they had an intensity value of exactly zero, a detection p>.01, or a bead count of<3. After these steps, probes that failed based on the above criteria in >5% of the samples were excluded from all samples (only probes with a success rate≥ 0.95 were retained). The methylation data were normalized with functional normalization (Fortin et al., 2014).

### DNA-methylation biomarker of biological aging

DNA methylation age acceleration measures of Hannum, Horvath, PhenoAge, and GrimAge were computed through the Horvath epigenetic age calculator (https://dnamage.genetics.ucla.edu/) and included the following outputs:

1. Hannum age acceleration—“AgeAccelerationResidualHannum” (AARH),
2. Intrinsic epigenetic age acceleration; the residual resulting from regressing the DNAm age estimate from Horvath on chronological age and blood cell count estimates.—“IEAA”,
3. PhenoAge acceleration—“AgeAccelPheno” (AAPheno),
4. GrimAge acceleration—“AgeAccelGrim” (AAGrim)

DunedinPACE was calculated based on code accessible on GitHub (https://github.com/danbelsky/DunedinPACE). For each of these epigenetic biomarkers, outliers were assessed and datapoints were removed when they were 3 times above or below the interquartile range (N_IEAA_ = 77, N_AARH_ = 57, N_AAPheno_ = 164, N_AAGrim_ = 118, N_DundinPACE_ = 59). The age acceleration values of the Hannum, Horvath, PhenoAge, and GrimAge biomarkers are on the same scale; they are expressed as the difference (in units of years) in estimated biological age and chronological. For instance, a value of 0 means that a person’s biological age is equal to their chronological age while values greater than 0 are interpreted as accelerated ageing and values smaller than 0 as decelerated aging. In contrast, the DunedinPACE algorithm does not estimate biological age (in units of years) but pace of ageing; a measure of how fast someone is ageing at the moment of sample collection. It is scaled such that a value of 1 corresponds to people who gain 1 year in biological age per year of chronological age. Values greater than 1 are interpreted as accelerated biological aging and values smaller than 1 are interpreted as decelerated biological aging.

### Life Events

NTR administered a Dutch life event scale (the Schokverwerkings Inventarisatie Lijst; Van der Velden et al., 1992; Middeldorp et al., 2008) in the 2009 survey. Participants were asked about traffic accident, violent assault, sexual assault, robbery, death of a spouse or child, serious illness or injury of self, spouse or child, job loss, and financial problems, with response categories “never experienced”, “less than 1 year ago”, “1–5 years ago” and “more than 5 years ago”. For divorce or break-up of a relationship equivalent to marriage, the response category was yes or no. All responses were recoded into binary variables indicating whether an individual had ever experienced each particular event prior to DNA collection, with ‘yes’ or ‘no’ as possible responses. The prevalence of each type of life event is given in Table 2. For our analyses, we also computed a sum score reflecting the total number of negative life events encountered by a participant, ranging from 0 to 9 as no participant had experienced all twelve life events.

**Table 2.** Prevalence of experiencing negative life events.

| Negative Life Event | Not experienced |  | Experienced |  |
| --- | --- | --- | --- | --- |
|  | N | % | N | % |
| Accident | 1413 | 72.2% | 395 | 20.2% |
| Violent Crime | 1687 | 86.2% | 121 | 6.2% |
| Sexual Crime | 1674 | 85.5% | 134 | 6.8% |
| Theft | 1282 | 65.5% | 526 | 26.9% |
| Death of Partner | 1780 | 90.9% | 28 | 1.4% |
| Death of Child | 1764 | 90.1% | 44 | 2.2% |
| Illness Partner | 1561 | 79.7% | 247 | 12.6% |
| Illness Child | 1616 | 82.5% | 192 | 9.8% |
| Illness Self | 1368 | 69.9% | 440 | 22.5% |
| Job loss | 1465 | 74.8% | 343 | 17.5% |
| Financial Problems | 1596 | 81.5% | 212 | 10.8% |
| End of Relationship | 1560 | 79.7% | 248 | 12.7% |
| **Sum Score** | **Mean** | **SD** | **Min** | **Max** |
| Negative Life Events | 1.621 | 1.619 | 0 | 9 |

Note: Those who experienced a life event can overlap. Sum Score depicts the total number of different events experienced.

### Covariates

#### Technical Covariates

To adjust for technical variation, array row and bisulfite plate (dummy-coding) were included as covariates.

#### Sex and Chronological Age

To adjust for possible sex and age effects both sex and chronological age (Kriegel et al, 2023), measured at time of blood draw, were included as covariates.

#### Lifestyle Covariates

BMI was computed based on reported weight and height (kg/m2) obtained at blood draw. Smoking status was classified as non-smoker (coded as 0), former-smoker (coded as 1), and current-smoker (coded as 2) obtained at blood draw.

#### Cell counts

After blood collection, cell counts were measured in fresh samples with the complete cell count method. The following subtypes of white blood cells were assessed: neutrophils, lymphocytes, monocytes, eosinophils, and basophils (Lin et al., 2017a; Lin et al., 2017b; Willemsen et al., 2010). Lymphocyte and neutrophil percentages were strongly negatively correlated (r=−0.93, p<0.001). Of these two white blood cell subtypes, the percentage of neutrophils showed the strongest correlation with DNA methylation levels (as evidenced by the correlation with PCs from the raw genome-wide methylation data). Basophil percentage showed little variation between subjects, with a large number of subjects (90,9%) having <1% of basophils. Therefore, the percentages of neutrophils, monocytes, and eosinophils were included as covariates in sensitivity analyses to adjust for inter-individual variation in white blood cell proportions.

### Data availability

The HumanMethylation450 BeadChip data from the NTR are available as part of the Biobank-based Integrative Omics Studies (BIOS) Consortium in the European Genome-phenome Archive (EGA), under the accession code EGAD00010000887. They are also available upon request via the BBMRI-NL BIOS consortium (https://www.bbmri.nl/acquisition-use-analyze/bios). All NTR data can be requested by researchers (https://ntr-data-request.psy.vu.nl/).

### Statistical analysis

Associations between experiencing negative life events and epigenetic age acceleration at the population level were tested in R in generalized estimating equation (GEE) models taking clustering within families into account. Initially, we assessed the impact of the total number of negative life events experienced by participants, creating a sum score ranging from 0 (no life events experienced) to 9. Subsequently, we analyzed the effect of individual life events. We first included all events in one model, each coded as 0 (not experienced) or 1 (experienced life event). Next, to address overlapping life events, we also conducted separate analyses for each life event, comparing participants who had not experienced any of the twelve negative life events to those who experienced specific ones. GEE models were fitted with the R package GEE, with the following specifications: Gaussian link function, 100 iterations, and the ‘exchangeable’ option to account for the correlation structure within families and within persons. For each of these analyses, we considered three models. In the primary model, we corrected for chronological age (as previously recommended by Krieger et al., 2023), sex, and technical covariates only. Additional models were fitted to examine whether the associations between negative life events and epigenetic aging might be driven by lifestyle or blood cell composition. In the second model, we additionally adjusted for smoking and BMI, and in the third model, we also included white blood cell proportions. We note that some (but not all) of the epigenetic biomarkers already correct for (estimated) white blood cell composition by design to derive a measure of age acceleration that is approximately independent of cellular composition and that one biomarker (Hannum – designed to represent a marker of aging of the immune system) explicitly incorporates estimated white blood cell composition in the calculation of biological age. If no significant association was found between an epigenetic biomarker and a life event in the primary model, then this association was not further analyzed in the second and third models.

After conducting the population-level analyses, we proceeded to examine the associations between the number of negative life events and epigenetic aging in discordant monozygotic twin pairs. Due to the small numbers of discordant monozygotic twin pairs for specific life events, such as loss of a child or partner, discordant twin pair analyses were not conducted separately for the specific life events. We therefore used the number of experienced life events to define discordance status within monozygotic twin pairs, where the twins who did not experience the same amount of life events were deemed discordant. We then performed a within-pair analysis, comparing epigenetic age acceleration between the twin who experienced the most negative life events and the twin who experienced the least negative life events. For the within-twin pair analyses, the same three models were considered, but the models did not correct for sex and age, as these are the same within MZ twin pairs. This analysis was done using fixed effects regression in R (Gonggrijp et al., 2023).

Correction for multiple testing was done by Bonferroni correction for the number of independent tests (α = 0.05/N independent tests). To estimate the number of independent variables in the correlation matrix of the dependent variables, we employed Matrix Spectral Decomposition (Nyholt, 2004) in R. A tetrachoric correlation matrix encompassing all 12 life events and a Spearman’s correlation matrix for the 5 epigenetic biomarkers was used as inputs (see Supplementary Table 1). We identified eight independent dimensions related to life events and three pertaining to the epigenetic biomarkers. This resulted in a cumulative total of 8*3=24 independent tests, leading to an adjusted alpha level of 0.002. Consequently, the corresponding confidence intervals in Figure 3 were adjusted to reflect this more stringent alpha level, thereby providing 99.8% confidence intervals for our estimates.

## Results

### Descriptive statistics

Table 2 presents the prevalence of various life events experienced by participants. Being the victim of theft emerged as the most common life event, experienced by 26.9% of the sample. This was closely followed by serious illness at 22.5%. The death of a partner and the death of a child had the lowest prevalence rates, 1.4% and 2.2% respectively. Only 548 (30.30%) participants did not experience any of the measured life events. Figure 2 presents a visualization of the various life events experienced by participants, highlighting the most common combinations of events. For instance, while a significant number of participants reported theft as an isolated event (N=109), others have experienced theft concurrently with other life events (e.g., 39 times with traffic accidents, 28 times with serious illness, and 28 times with job loss). Figure 2 includes only those groups or combinations of events where at least 10 individuals are represented. Supplementary Figure 1 presents a more exhaustive visualization that includes all reported combinations of life events, regardless of their frequency. The intricate web of intersections in the matrix section of both Figure 2 and Supplementary Figure 1 indicates that a majority of participants experienced more than one negative life event (N=790), rather than a single isolated incident. The live events sum score, representing the total number of different life events encountered, had an average of 1.62 (SD 1.62).

**Figure 2.**
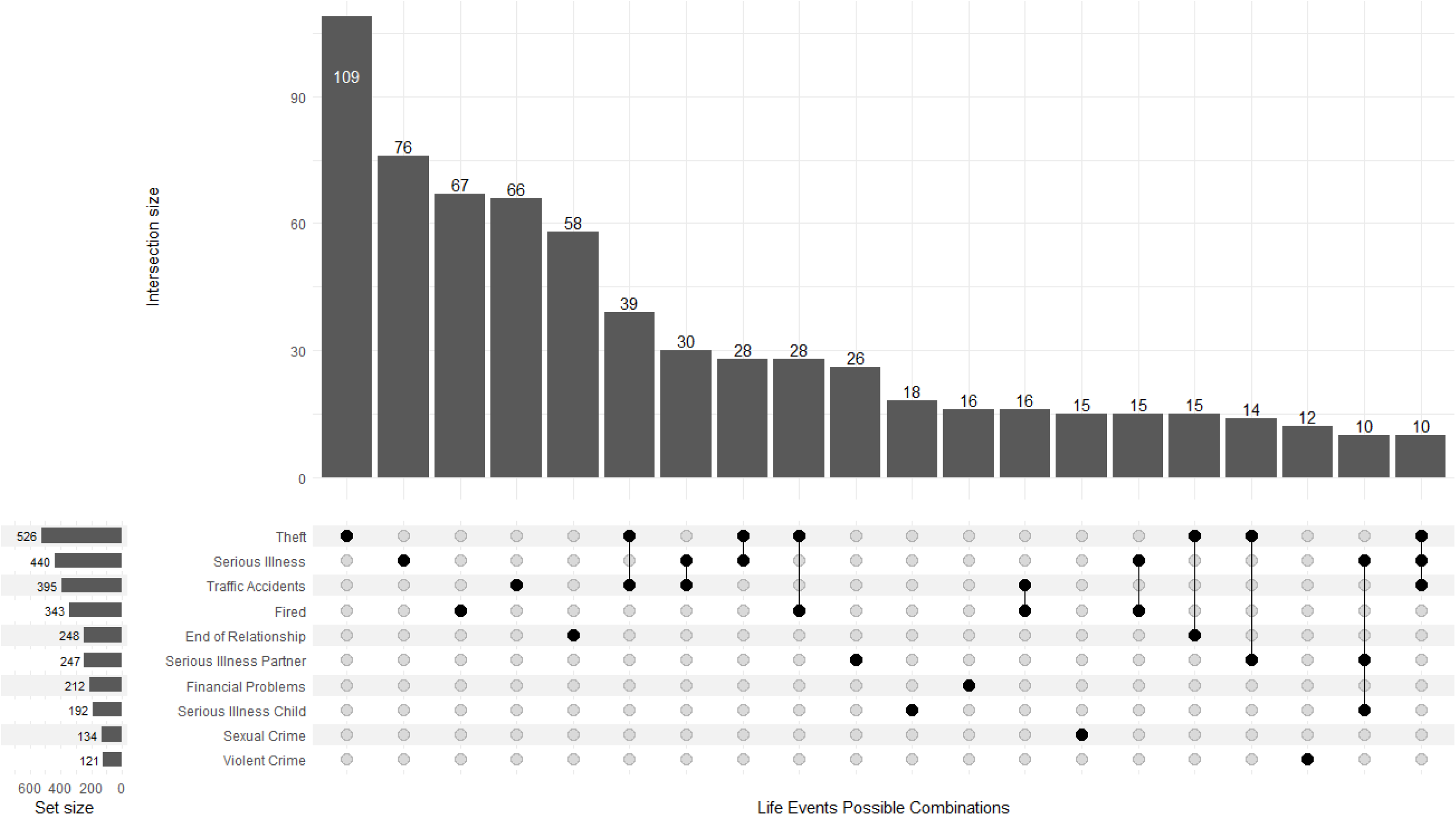
Visualization of Singular and Overlapping Life Events Experienced. Note: Horizontal bars on the left side (set size) denote the total number of participants who experienced each specific life event, while the vertical bars at the top showcase the number of pants who experienced combinations of events. The matrix in the middle indicates which events are included in each combination. Only combinations with 10 participants or more are d in the graph.

**Figure 3.**
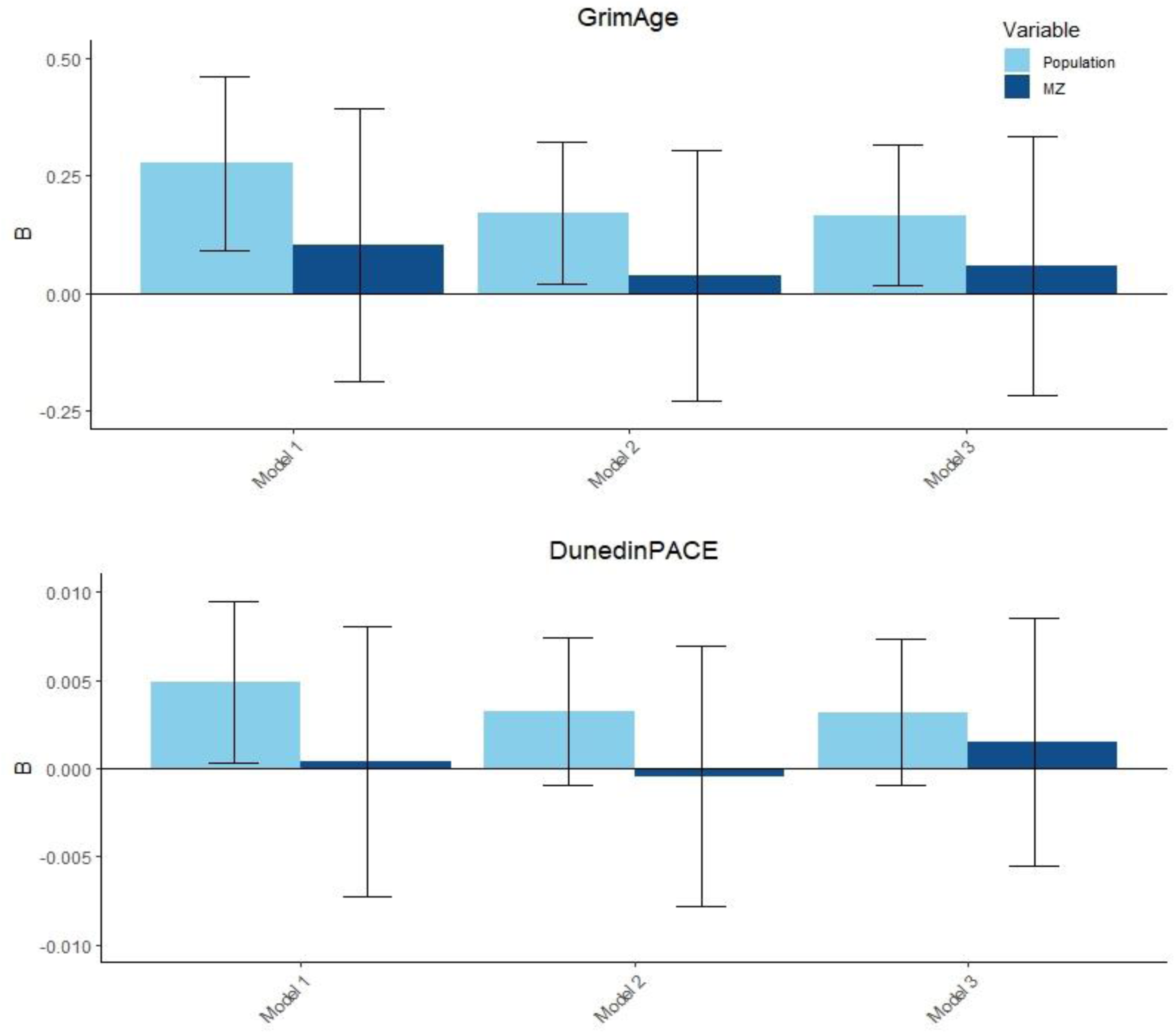
Associations of GrimAge and DunedinPACE with Total Number of Life Events: Population vs. Discordant MZ Twin Analyses for the Different Models. Note: Model 1 corrects for age, sex, and technical covariates. Model 2 additionally adjusts for BMA and smoking, and Model 3 additionally adjusts for white blood cell count. 99.8% CI (p<.002).

Table 3 displays the descriptive statistics for the epigenetic biomarkers, white blood cell counts, BMI, and smoking status. The mean age of the participants was 37.69 (SD = 13.23), and the majority of the participants were female (70.30%).

**Table 3.**
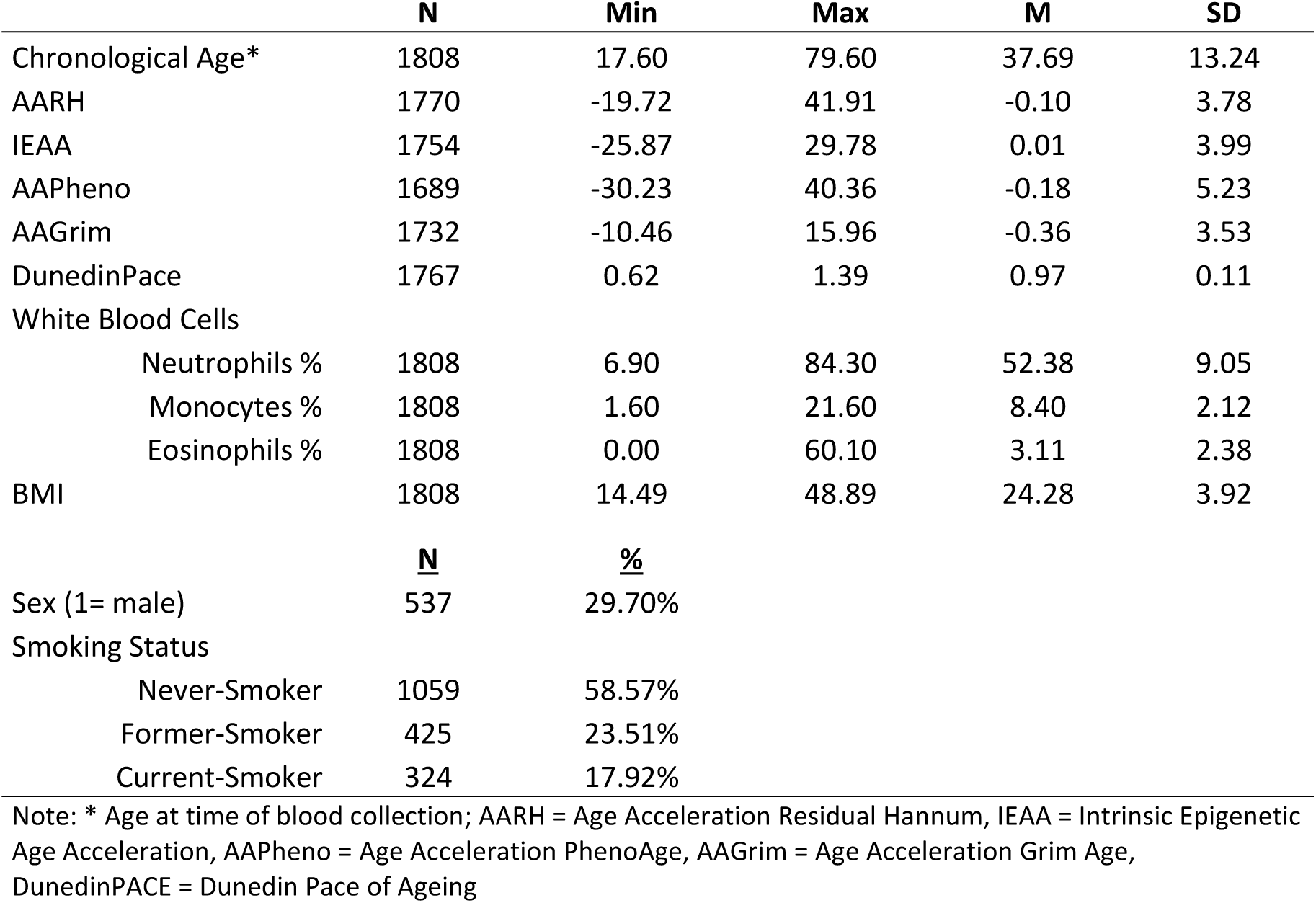
Descriptive statistics of the epigenetic biomarkers, chronological age, white blood cell counts, sex, BMI, and smoking status.

### Population Analyses

Prior to the population analyses, a GEE was conducted to look at the associations between the epigenetic biomarkers and the covariates as well as the life events and the covariates, see Supplementary Tables 2 and 3. Almost all covariates showed a significant relationship with the 5 epigenetic biomarkers. For example, a higher BMI was associated with epigenetic age acceleration for the PhenoAge, GrimAge, and DunedinPACE biomarkers. Sex differences were also observed for most, except for PhenoAge, with men showing an acceleration compared to women in the epigenetic biomarkers Hannum, Horvath, and GrimAge and a deceleration in the DunedinPACE biomarker. Therefore, these findings underscore the importance of considering factors like BMI and sex in the analysis of epigenetic aging.

Results of the first analyses, in which associations were adjusted for chronological age, sex, and technical covariates, are outlined in Table 4. The Hannum, Horvath, and PhenoAge did not exhibit significant associations with the total number of negative life events, or any of the specific life events examined. These biomarkers were therefore excluded from further analysis. In contrast, the upper part of Table 4 shows that the GrimAge and DunedinPACE biomarkers demonstrated a notable increase in epigenetic aging related to the total number of experienced life events B_GrimAA_ = 0.276, p= 3.79E-06 and B_DunedinPACE_ = 0.005, p= 0.001). The regression coefficient of 0.276 for GrimAge implies that each additional life event experienced is associated with an acceleration of biological aging by almost 0.3 years. Therefore, if an individual has experienced four life events, this corresponds to 0.276 * 4 = 1.104 additional years of biological aging in comparison to someone who has not experienced any such events. The DunedinPACE biomarker is a pace of aging biomarker, and thus the regression coefficient of 0.005 indicates that each additional life event is associated with an accelerated aging by 0.005 years per calendar year. Therefore, if an individual experiences four life events, the accumulated effect would be an extra 0.02 years (0.005 *4) of biological aging per year, which corresponds to an increased pace of aging of about 1 week per year. The lower part of Table 4 shows the results of the subsequent analyses in which the associations with all negative life events were estimated simultaneously. These analyses revealed that only financial adversities were linked to an age acceleration of approximately 1.2 years in GrimAge (B = 1.215, p = 1.09E-4) when all life events were considered concurrently.

**Table 4.**
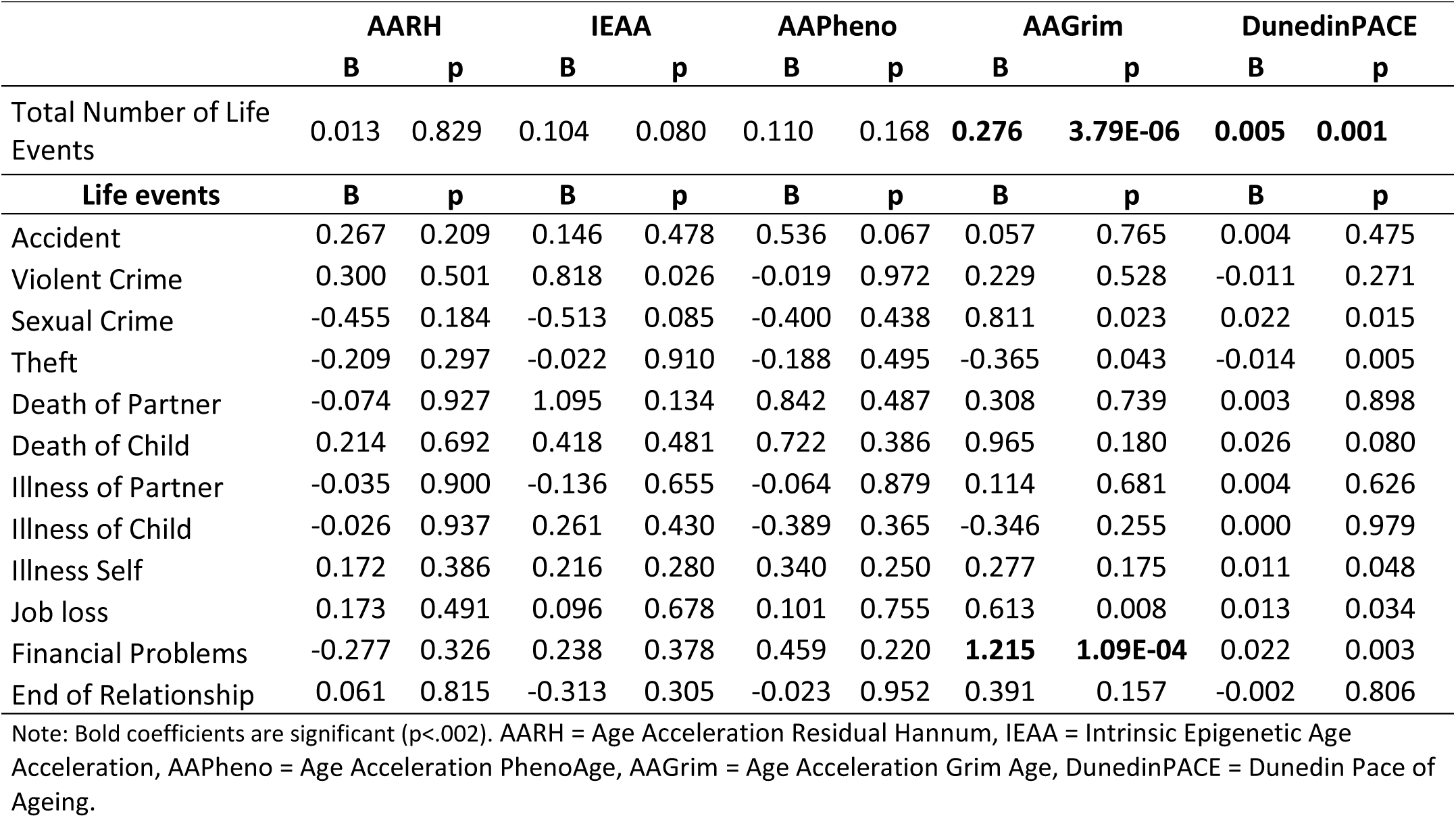
Primary GEE Model: Associations Between Epigenetic Aging and All Negative Life Events Estimated Simultaneously, Adjusted for Age, Sex, and Technical Covariates.

We also conducted additional GEE analyses in which respondents who experienced each individual life event were compared to a control group of respondents who reported no life events. This approach was intended to isolate the impact of each specific life event. Results are shown in table 5. These analyses reinforced the association between financial adversities and an acceleration in GrimAge (B = 1.536, p = 1.26E-5) and further identified significant associations for sexual crimes (B = 1.544 p = 1.14E-04) and job loss (B = 0.868, p = 0.001). The DunedinPACE biomarker similarly showed acceleration for sexual crimes (B = 0.034, p = 0.001) and financial problems (B = 0.032, p = 2.76E-04). Both financial adversities and experiencing sexual crimes were thus associated with approximately 1.5 years of age acceleration for the GrimAge biomarker, and an increased pace of aging of 0.03 years, roughly 12 days, per calendar year to DunedinPACE.

**Table 5.**
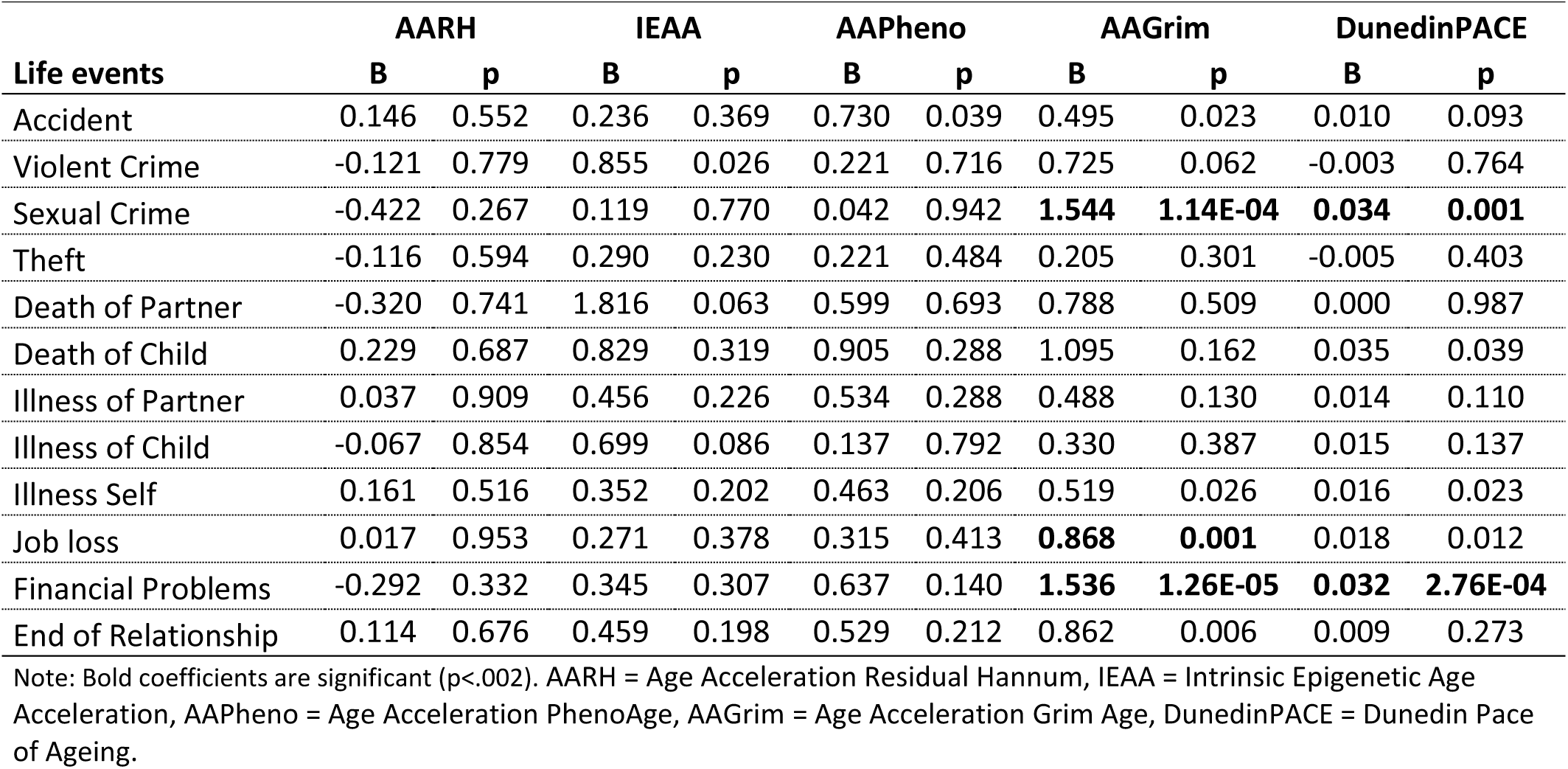
Primary GEE Model: Associations Between Epigenetic Aging and Negative Life Events Estimated Separately, Adjusted for Age, Sex, and Technical Covariates.

| Life events | AARH |  | IEAA |  | AAPhen |  | AAGrim |  | DunedinPACE |  |
| --- | --- | --- | --- | --- | --- | --- | --- | --- | --- | --- |
|  | B | p | B | p | B | p | B | p | B | p |
| Accident | 0.146 | 0.552 | 0.236 | 0.369 | 0.730 | 0.039 | 0.495 | 0.023 | 0.010 | 0.093 |
| Violent Crime | -0.121 | 0.779 | 0.855 | 0.026 | 0.221 | 0.716 | 0.725 | 0.062 | -0.003 | 0.764 |
| Sexual Crime | -0.422 | 0.267 | 0.119 | 0.770 | 0.042 | 0.942 | **1.544** | **1.14E-04** | **0.034** | **0.001** |
| Theft | -0.116 | 0.594 | 0.290 | 0.230 | 0.221 | 0.484 | 0.205 | 0.301 | -0.005 | 0.403 |
| Death of Partner | -0.320 | 0.741 | 1.816 | 0.063 | 0.599 | 0.693 | 0.788 | 0.509 | 0.000 | 0.987 |
| Death of Child | 0.229 | 0.687 | 0.829 | 0.319 | 0.905 | 0.288 | 1.095 | 0.162 | 0.035 | 0.039 |
| Illness of Partner | 0.037 | 0.909 | 0.456 | 0.226 | 0.534 | 0.288 | 0.488 | 0.130 | 0.014 | 0.110 |
| Illness of Child | -0.067 | 0.854 | 0.699 | 0.086 | 0.137 | 0.792 | 0.330 | 0.387 | 0.015 | 0.137 |
| Illness Self | 0.161 | 0.516 | 0.352 | 0.202 | 0.463 | 0.206 | 0.519 | 0.026 | 0.016 | 0.023 |
| Job loss | 0.017 | 0.953 | 0.271 | 0.378 | 0.315 | 0.413 | **0.868** | **0.001** | 0.018 | 0.012 |
| Financial Problems | -0.292 | 0.332 | 0.345 | 0.307 | 0.637 | 0.140 | **1.536** | **1.26E-05** | **0.032** | **2.76E-04** |
| End of Relationship | 0.114 | 0.676 | 0.459 | 0.198 | 0.529 | 0.212 | 0.862 | 0.006 | 0.009 | 0.273 |

Note: Bold coefficients are significant ( $p < .002$ ). AARH = Age Acceleration Residual Hannum, IEAA = Intrinsic Epigenetic Age Acceleration, AAPhen = Age Acceleration PhenoAge, AAGrim = Age Acceleration Grim Age, DunedinPACE = Dunedin Pace of Ageing.

Additional sensitivity analyses were performed for significant associations. In the second model, adjusted for BMI and smoking, several associations from the basic model were no longer statistically significant, and effect sizes were reduced, although the direction of effects remained the same, as detailed in Supplementary Table 4. A notable change was observed with the DunedinPACE biomarker. After accounting for BMI and smoking, all previously significant associations related to DunedinPACE lost their statistical significance, and effect sizes were reduced by nearly half. This suggests that lifestyle factors such as BMI and smoking significantly impact the associations observed with this particular biomarker. In the case of the GrimAge biomarker, after incorporating BMI and smoking as covariates, the age acceleration link with experiencing sexual crimes and job loss no longer held statistical significance, and their effect sizes were reduced by half after the additional correction for lifestyle factors. The total number of life events continued to show a significant relationship with age acceleration for GrimAge, but the effect size was reduced by 38.04% with each experienced life event showing an increase of 0.171 years (p = 4.24E-04). Additionally, when all life events were considered in the same model, financial problems showed no significant age accelerations, and the effect size was reduced by half. However, when comparing the effects of life events separately with those who did not experience any life events, experiencing financial problems continued to show an acceleration regarding the GrimAge biomarker (B= 0.885, p = 0.002).

Table 6 presents the outcomes of the third model, which took into account chronological age, sex, technical covariates, lifestyle factors, and white blood cell counts when estimating the associations with all life events simultaneously. Additionally, Table 7 presents the outcome of the third model with the life events considered separately. This comprehensive approach led to the attenuation of several associations observed in the basic model. However, the GrimAge biomarker continued to demonstrate significant associations, indicative of its robustness across various model specifications. Table 6 shows that the total number of life effects was associated with an increase in epigenetic aging. Specifically, each experienced life event was linked to an age acceleration of 0.166 years (p = 0.001), thus for those who experienced 9 life events, this was linked to an increase of 0.166 * 9 = 1.5 years. In terms of specific life events, the patterns observed in the second model persisted. Notably, the results in Table 7 show that financial difficulties continued to exhibit the strongest association with age acceleration, contributing to an increase of 0.855 years in epigenetic age in comparison to those who did not experience any of the life events.

**Table 6.** Third GEE Model: Associations Between Epigenetic Aging and All Negative Life Events Estimated Simultaneously, adjusted for Lifestyle Factors and White Blood Cell Counts Age, Sex, and Technical Covariates.

|  | GrimAge |  | DunedinPACE |  |
| --- | --- | --- | --- | --- |
|  | B | p | B | p |
| Total number of Life Events | **0.166** | **0.001** | 0.003 | 0.019 |
| **Life events** | **B** | **p** | **B** | **p** |
| Financial Problems* | 0.659 | 0.008 |  |  |

Note: Bold coefficients are significant ( $p < .002$ ), \* all other 11 life events are included in the model.

**Table 7.** Third GEE Model: Associations Between Epigenetic Aging and All Negative Life Events Estimated Separately, adjusted for Lifestyle Factors and White Blood Cell Counts Age, Sex, and Technical Covariates.

| Life events | GrimAge |  | DunedinPACE |  |
| --- | --- | --- | --- | --- |
|  | B | p | B | p |
| Sexual Crime | 0.738 | 0.021 | 0.019 | 0.025 |
| Job loss | 0.384 | 0.070 |  |  |
| Financial Problems | **0.855** | **0.002** | 0.018 | 0.014 |

Note: Bold coefficients are significant ( $p < .002$ )

### Discordant Twin Pair Analyses

Supplementary Table 5 shows the numbers of concordant and discordant MZ twin pairs for the number of life events. In the group of 421 MZ twin pairs, we observed 147 (34.92%) pairs that were concordant for the total number of life events and 274 (65.08%) pairs that were discordant. Following the population analyses, discordant MZ twin pair analyses were conducted for GrimAge and DunedinPACE and the total number of life events experienced in 274 MZ discordant pairs. Full results of the discordant twin analyses can be found in Supplementary Table 6. The discordant twin analysis showed associations between the total number of life events and the biomarkers GrimAge or DunedinPACE that were indicative of age acceleration, except for a small deceleration in Model 2 regarding the DunedinPACE biomarker. However, none of these associations were significant. Figure 4 illustrates the comparison between the population analyses and the discordant twin analyses for the GrimAge and DunedinPACE biomarkers, showcasing results with the adjusted 99.8% confidence intervals due to the adjusted alpha level of 0.002. It clearly shows that the associations observed in the population sample in all models were reduced by more than half in the MZ twin analyses, suggesting genetic and/or shared environmental confounding.

## Discussion

Our study examined the impact of a broad spectrum of negative life events on epigenetic aging by analyzing five epigenetic biomarkers, leveraging data from the Netherlands Twin Register. This approach allowed us to perform both between-individual and within-twin comparisons, providing a nuanced view of how negative life events may affect the aging process. The results of our first analyses showed that the total number of experienced life events is associated with an acceleration of biological aging for the GrimAge and DunedinPACE biomarkers. Further analysis refined these findings by separately evaluating the influence of distinct life events. Initially, by estimating associations for all life events simultaneously in a single model, financial problems showed a clear age acceleration for the GrimAge biomarker. To isolate the effects of specific life events, we then compared respondents who experienced each life event against a control group that experienced no life events. This separate analysis highlighted not only the pronounced effect of financial problems but also revealed that sexual crimes were significantly associated with an increase of approximately 1.5 years of age acceleration for the GrimAge biomarker and job loss with 0.86 years. The DunedinPACE biomarker paralleled these findings, showing an increased pace of aging of roughly 12 days per chronological year for individuals who encountered sexual crimes and financial problems. These insights show that not only the quantity but also the nature of life events are associated with epigenetic aging. Our findings resonate with and extend previous research that reported links between epigenetic biomarkers and childhood adversities, socioeconomic challenges, and PTSD (Joshi, Lin & Raine, 2023; Raffington et al., 2021; Katrinli et al., 2020; Yang et al., 2020). Our results extend these associations to a wider array of life events, with the most profound impacts observed in the context of financial strain and sexual crimes.

Diving deeper into our results, we incorporated additional covariates, specifically BMI, smoking habits, and white blood cell counts, to examine their influence on our previously observed associations. Our analyses revealed that the incorporation of BMI, smoking habits, and white blood cell counts significantly modified the associations between life events and epigenetic aging. Notably, in the second model, which accounted for BMI and smoking, several associations identified in the basic model lost their statistical significance, and the effect sizes were generally reduced. This was particularly evident with the DunedinPACE biomarker, where all previously significant associations became non-significant upon adjusting for these lifestyle factors. This shift underscores the potential confounding effects of these covariates, suggesting that the observed acceleration in biological aging, as measured by biomarkers like GrimAge and DunedinPACE, might be more intricately linked to broader lifestyle factors and physiological changes rather than directly attributable to life events themselves. The diminished significance of our findings post-adjustment indicates a complex interplay between life events, lifestyle choices, and biological factors. For instance, it raises the possibility that lifestyle factors such as smoking or variations in BMI, potentially altered in response to life stressors, could be primary drivers of epigenetic changes. Similarly, fluctuations in white blood cell counts, indicative of various physiological responses, might mediate the relationship between life events and biological aging. This complexity necessitates a cautious interpretation of the direct impact of life events on epigenetic aging. The most comprehensive model, which included additional adjustments for white blood cell counts alongside lifestyle factors, reaffirmed the associations with the GrimAge biomarker, and both the total number of life events experienced and financial problems, albeit with reduced effect sizes. This persistence suggests a potentially robust link between specific stressors and epigenetic aging, warranting further investigation. These observations are in line with previous research which found that that PTSD was associated with GrimAge acceleration (Yang et al., 2020, and Katrinli et al., 2020).

Our study further delved into the age acceleration findings from the population analyses by conducting discordant MZ twin analyses. By examining MZ twin pairs with differing life event experiences, we controlled for shared genetic and environmental factors, providing a more unobstructed view of the relationship between life adversities and biological aging. In our sample of 421 MZ pairs, we found that 274 pairs were discordant and 147 concordant for the number of life events experienced. The high concordance in MZ pairs is consistent with genetic modeling of the life event data (Middeldorp et al. 2005) which indicates that genetic and common environmental factors play a substantial role in the individual differences in the experience of life events. This finding aligns with the notion that genetic predispositions may influence the likelihood of encountering or perceiving certain life events. However, the limited number of discordant MZ twin pairs regarding specific life events implies a lower statistical power. Consequently, we chose to focus our analysis on the total number of life events, rather than specific ones, as the variability within discordant pairs was insufficient to robustly assess the impact of specific life events. Intriguingly, when looking at the association between the total number of life events experienced and age acceleration in terms of GrimAge and DunedinPACE, the effect sizes of the population were reduced by more than half in the discordant twin analyses. This finding suggests that the observed acceleration at the population level in biological aging requires a more complex and nuanced explanation than that of a direct causal effect by life events. Rather than expressing a causal effect, the found associations are more likely confounded by shared family factors, both genetic and environmental. However, it is also possible that the impact of a life event experienced by one twin could also extend to the co-twin, particularly in cases of more severe incidents like the loss of a child and sexual crimes. The shared environment and emotional bonds between twins mean that a traumatic event affecting one twin could also have psychological, behavioral, or even biological repercussions for the other. Alternatively, financial and emotional support from a twin may reduce the impact of a life event. These inter-twin influences might lead to an underestimation of the effects in the MZ twin model. This phenomenon highlights a potential limitation in the MZ twin model, as it assumes independent exposure to life events for each twin. To further unravel the complexities observed in this study, future research should enhance discordant twin pair analyses, potentially with larger sample sizes, to assess the persistence of associations when adjusting for genetic and environmental influences. This approach will be crucial in clarifying the extent to which genetic and shared environmental factors may confound the relationship between life events and epigenetic aging

Despite the strengths of our study, it is not without its limitations. Firstly, while our sample size exceeded that of previous studies on negative life events and age acceleration, the specific nature of some events, such as the death of a child or partner or experiencing sexual crimes, meant that only a small number of individuals were affected. This limitation reduced our statistical power to detect minor effects and constrained our ability to thoroughly examine these specific events in the discordant twin samples, thereby limiting our capacity to draw definitive conclusions about these particular life events.

Secondly, the subjective nature of how individuals perceive and report life events presents another challenge (Kessler & Wethington, 1991; Van de Mortel, 2008). For instance, the significance of an event like theft can vary greatly; one person may not consider the theft of a bicycle significant enough to report, while another might. Although one might also argue that it is the subjective experience that matters rather than the objective severity of the event, this subjectivity still poses a challenge in standardizing and comparing experiences across individuals. Furthermore, there is substantial variability in the severity of incidents within the same category. For instance, a home burglary is generally more severe than bicycle theft and severe assault is more impactful than a mere threat. This variation in severity, coupled with the subjective experience of respondents, makes it complex to accurately categorize and assess the impact of life events. Future research, including longitudinal studies, larger sample sizes, and more additional information about event severity, and the support that people receive after having experienced a life event, could help to unravel these complex relationships.

In conclusion, our study has shed light on the intricate interplay between life events and epigenetic aging, highlighting the profound impact of experiencing life events on the biological aging processes. While our analyses reveal that lifestyle factors and white blood cell counts can partly explain the association between experiencing negative life events and aging, our findings highlight the significant impact of financial difficulties and experiencing multiple life events on biological aging, even after accounting for these variables. This suggests that policy initiatives aimed at reducing financial hardships among citizens could also have positive implications for their health. By focusing on lessening financial stress, these policies might contribute to mitigating the accelerated aging associated with such stressors. The discordant monozygotic twin analysis adds a layer of complexity to our understanding as they suggested that familial factors may confound the observed associations. This outcome highlights the need for caution in interpreting these relationships between experiencing negative life events and epigenetic aging and underscores the importance of further research in this area. This study not only reinforces the importance of considering a broad spectrum of life events in epigenetic aging research but also calls for more nuanced, longitudinal studies that can address the limitations we identified, such as the varying interpretations of life event severity and the need for larger sample sizes for rare events. This work reinforces the importance of considering both genetic and environmental factors in the study of epigenetics and aging.

## Funding and acknowledgements

This study was supported by the Victim Support Fund (Fonds Slachtofferhulp), The Netherlands. NTR: Data collection in the Netherlands Twin Register was funded by the Netherlands Organization for Scientific Research (NWO), grant numbers 31160008, 904-61-193 and 985-10-002. The Netherlands Organization for Scientific Research (NWO): Biobanking and Biomolecular Research Infrastructure (BBMRI-NL, NWO 184.033.111) and the BBRMI-NL-financed BIOS Consortium (NWO 184.021.007), NWO Large Scale infrastructures X-Omics (184.034.019), Genotype/phenotype database for behaviour genetic and genetic epidemiological studies (ZonMw Middelgroot 911-09-032); Netherlands Twin Registry Repository: researching the interplay between genome and environment (NWO-Groot 480-15-001/674); epigenetic data were generated at the Human Genomics Facility (HuGe-F) at ErasmusMC Rotterdam.

We warmly thank all twin families for their participation.

## References

Bell C.G., Lowe R, Adams P.D., Baccarelli A.A., Beck S, Bell J.T., et al. DNA methylation aging biomarkers: challenges and recommendations. Genome Biol. 2019[cited 2019 Dec 2];20(1):249

Belsky, D. W., Caspi, A., Corcoran, D. L., Sugden, K., Poulton, R., Arseneault, L., … & Moffitt, T. E. (2022). DunedinPACE, a DNA methylation biomarker of the pace of aging.

Boks, M. P., & Mierlo, H. C. (2015). van, Rutten BPF, TRDJ Radstake, De Witte L, Geuze E et al. Longitudinal changes of telomere length and epigenetic age related to traumatic stress and post-traumatic stress disorder. Psychoneuroendocrinology, 51, 506–512.

Brown, D. W., Anda, R. F., Tiemeier, H., Felitti, V. J., Edwards, V. J., Croft, J. B., & Giles, W. H. (2009). Adverse childhood experiences and the risk of premature mortality. American journal of preventive medicine, 37(5), 389–396.

Dag, Y. N., Mehlig, K., Rosengren, A., Lissner, L., & Rosvall, M. (2020). Negative emotional states and negative life events: Consequences for cardiovascular health in a general population. Journal of Psychosomatic Research, 129, 109888.

Distel M.A., Middeldorp C.M., Trull T.J., Derom C.A., Willemsen G, Boomsma D.I. (2011). Life events and borderline personality features: the influence of gene-environment interaction and gene-environment correlation. Psychol Med. 41(4):849–60.

Fortin J.P., Labbe A, Lemire M, Zanke B.W., Hudson T.J., Fertig E.J., et al. (2014) Functional normalization of 450k methylation array data improves replication in large cancer studies. Genome Biol 15, 503

Gesell, A. (1942). The method of co-twin control. Science, 95, 446–448.

Gonggrijp, B. M., van de Weijer, S. G., Bijleveld, C. C., van Dongen, J., & Boomsma, D. I. (2023). The Co-Twin Control Design: Implementation and Methodological Considerations. Twin Research and Human Genetics, 1-8.

Hannum, G., et al., Genome-wide methylation profiles reveal quantitative views of human aging rates. Mol Cell, 2013. 49(2): p. 359–367.

Horvath, S. (2013) DNA methylation age of human tissues and cell types. Genome Biol 14, doi:10.1186/gb-2013-14-10-r115.

de Jong, R. (2022). Long-term consequences of child sexual abuse: Adult role fulfillment of child sexual abuse victims.

Katrinli, S., Stevens, J., Wani, A. H., Lori, A., Kilaru, V., van Rooij, S. J., … & Smith, A. K. (2020). Evaluating the impact of trauma and PTSD on epigenetic prediction of lifespan and neural integrity. Neuropsychopharmacology, 45(10), 1609–1616.

Kessler R.C. & Wethington E. The reliability of life event reports in a community survey. Psychological Medicine. 1991;21(3):723–738.

Kim, K., Yaffe, K., Rehkopf, D. H., Zheng, Y., Nannini, D. R., Perak, A. M., … & Hou, L. (2023). Association of adverse childhood experiences with accelerated epigenetic aging in midlife. JAMA network open, 6(6), e2317987–e2317987.

Krieger, N., Chen, J. T., Testa, C., Diez Roux, A., Tilling, K., Watkins, S., … & Relton, C. (2023). Use of correct and incorrect methods of accounting for age in studies of epigenetic accelerated aging: implications and recommendations for best practices. American Journal of Epidemiology, 192(5), 800–811.

Levine, M. E., Lu, A. T., Bennett, D. A., & Horvath, S. (2015). Epigenetic age of the pre-frontal cortex is associated with neuritic plaques, amyloid load, and Alzheimer’s disease related cognitive functioning. Aging (Albany NY), 7(12), 1198.

Ligthart, L., van Beijsterveldt, C. E. M., Kevenaar, S. T., De Zeeuw, E., van Bergen, E., Bruins, S, Pool, R., Helmer, Q, van Dongen, J., Hottenga, J. J. van’t Ent, D., Dolan, C. V., Davies, G. E., Ehli, E. A., Bartels, M., Willemsen, G., de Geus, E. J.C., & Boomsma, D. I. (2019). The Netherlands Twin Register: Longitudinal Research Based on Twin and Twin-Family Designs. Twin Research and Human Genetics, 22, 623–636.

Lin, B. D., Willemsen, G., Fedko, I. O., Jansen, R., Penninx, B., De Geus, E., … & Boomsma, D. I. (2017a). Heritability and gwas studies for monocyte–lymphocyte ratio. Twin Research and Human Genetics, 20(2), 97–107.

Lin, B. D., Carnero-Montoro, E., Bell, J. T., Boomsma, D. I., De Geus, E. J., Jansen, R., … & Hottenga, J. J. (2017b). 2SNP heritability and effects of genetic variants for neutrophil-to-lymphocyte and platelet-to-lymphocyte ratio. Journal of human genetics, 62(11), 979–988.

Marioni, R. E., Shah, S., McRae, A. F., Chen, B. H., Colicino, E., Harris, S. E., … & Deary, I. J. (2015a). DNA methylation age of blood predicts all-cause mortality in later life. Genome biology, 16(1), 1–12.

Marioni, R. E., Shah, S., McRae, A. F., Ritchie, S. J., Muniz-Terrera, G., Harris, S. E., … & Deary, I. J. (2015b). The epigenetic biomarker is correlated with physical and cognitive fitness in the Lothian Birth Cohort 1936. International journal of epidemiology, 44(4), 1388–1396.

Middeldorp, C. M., Cath, D. C., Beem, A. L., Willemsen, G., & Boomsma, D. I. (2008). Life events, anxious depression and personality: a prospective and genetic study. Psychological medicine, 38(11), 1557–1565.

Monaghan, P., & Haussmann, M. F. (2015). The positive and negative consequences of stressors during early life. Early Human Development, 91(11), 643–647.

Nyholt D.R. A simple correction for multiple testing for single-nucleotide polymorphisms in linkage disequilibrium with each other. Am J Hum Genet. 2004;74(4):765–9

Oblak, L., van der Zaag, J., Higgins-Chen, A. T., Levine, M. E., & Boks, M. P. (2021). A systematic review of biological, social and environmental factors associated with epigenetic biomarker acceleration. Ageing research reviews, 69, 101348.

Patton, G. C., Coffey, C., Posterino, M., Carlin, J. B., & Bowes, G. (2003). Life events and early onset depression: cause or consequence?. Psychological medicine, 33(7), 1203–1210.

Pridemore, W. A., & Berg, M. T. (2017). What is past is prologue: A population-based case-control study of repeat victimization, premature mortality, and homicide. Aggressive behavior, 43(2), 176–189.

Joshi, D., Gonzalez, A., Lin, D., & Raina, P. (2023). The association between adverse childhood experiences and epigenetic age acceleration in the Canadian longitudinal study on aging (CLSA). *Aging Cell*, e13779.

Raffington, L., Belsky, D. W., Kothari, M., Malanchini, M., Tucker-Drob, E. M., & Harden, K. P. (2021). Socioeconomic disadvantage and the pace of biological aging in children. Pediatrics, 147(6).

Raffington, L., & Belsky, D. W. (2022). Integrating DNA methylation measures of biological aging into social determinants of health research. Current Environmental Health Reports, 9(2), 196–210.

Sumner, J. A., Gao, X., Gambazza, S., Dye, C. K., Colich, N. L., Baccarelli, A. A., … & McLaughlin, K. A. (2023). Stressful life events and accelerated biological aging over time in youths. Psychoneuroendocrinology, 151, 106058.

Van de Mortel, T. F. (2008). Faking it: social desirability response bias in self-report research. Australian Journal of Advanced Nursing, The, 25(4), 40–48.

Van der Velden, P. G., Contino, C., van de Ven, P., & Das, M. (2021). The use of professional help and predictors of unmet needs for dealing with mental health to legal problems among victims of violence, accidents, theft and threat, and nonvictims in the general population. PLoS one, 16(11), e0259346.

Van Dongen J, Nivard M.G., Willemsen G, Hottenga J.J., Helmer Q, Dolan C.V., et al. (2016) Genetic and environmental influences interact with age and sex in shaping the human methylome. Nat Commun 7;7:11115

Van Iterson M, Tobi E.W., Slieker R.C., Den Hollander W, Luijk R, Slagboom P.E., et al. (2014). MethylAid: Visual and interactive quality control of large Illumina 450k datasets. Bioinformatics.

Wang Z, Hui Q, Goldberg J, Smith N, Kaseer B, Murrah N, Levantsevych OM, Shallenberger L, Diggers E, Bremner JD, Vaccarino V, Sun YV. (2022) Association Between Posttraumatic Stress Disorder and Epigenetic Age Acceleration in a Sample of Twins. Psychosom Med. 01;84(2):151–158.

Willemsen G, de Geus E.J., Bartels M, van Beijsterveldt C.E., Brooks A.I., Estourgie-van Burk G.F. et al. (2010) The Netherlands Twin Register biobank: a resource for genetic epidemiological studies. Twin Res Hum Genet 13: 231–245.

Wolf, E., Logue, M., Morrison, F., Wilcox, E., Stone, A., Schichman, S., … Miller, M. (2019). Posttraumatic psychopathology and the pace of the epigenetic biomarker: A longitudinal investigation. Psychological Medicine, 49(5), 791–800.

Yang, G.W.Y. Wu, J.E. Verhoeven, A. Gautam, V.I. Reus, J.I. Kang, et al. (2020) A DNA methylation biomarker associated with age-related illnesses and mortality is accelerated in men with combat PTSD Mol. Psychiatry

Young, M., & Schieman, S. (2012). When hard times take a toll: The distressing consequences of economic hardship and life events within the family-work interface. Journal of health and social behavior, 53(1), 84–98.

