## Supplementary figures and images for "Negative Life Events and Epigenetic Ageing: a Study in the Netherlands Twin Register"

### Supplemental Figure 1

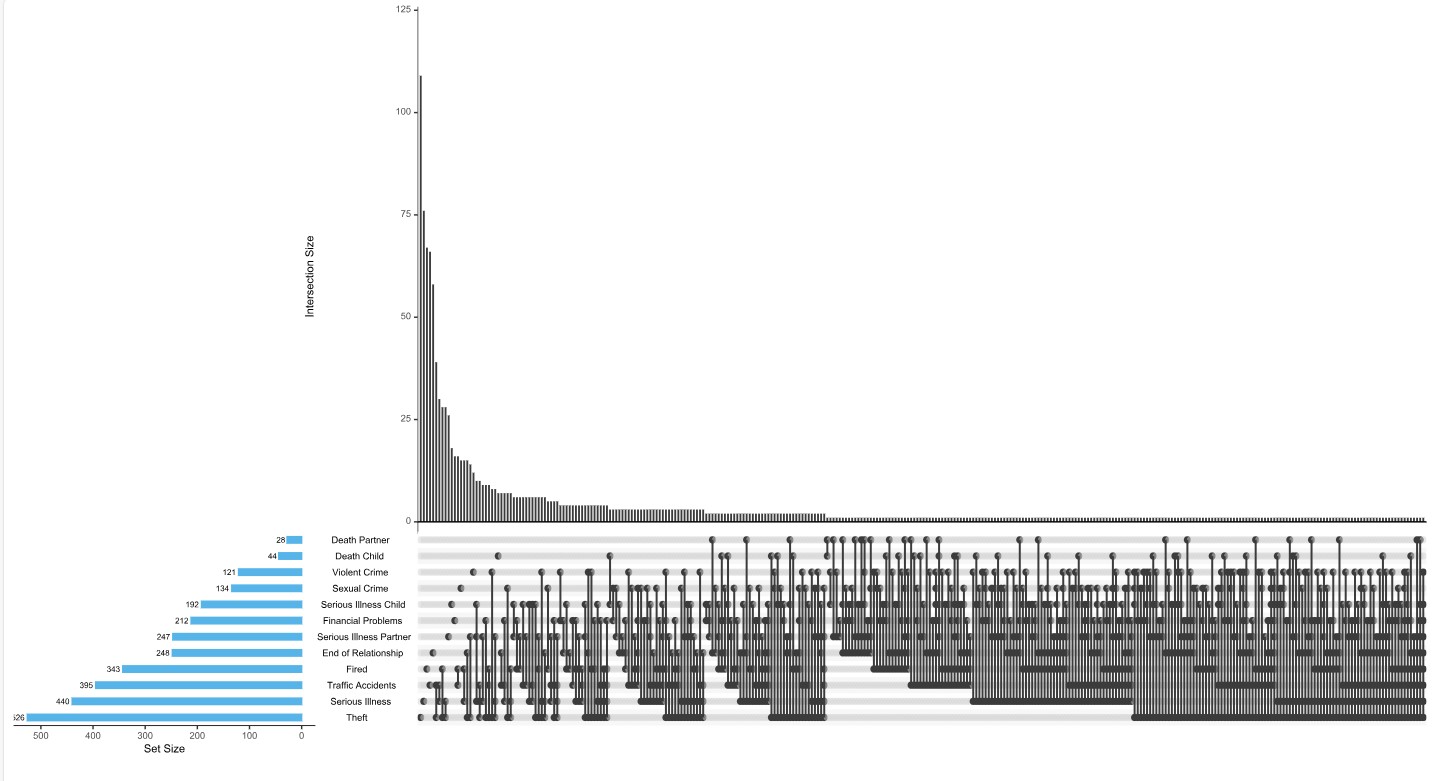
